# Contemporary evolution of a Lepidopteran species, *Heliothis virescens*, in response to modern agricultural practices

**DOI:** 10.1101/103382

**Authors:** Megan L Fritz, Alexandra M DeYonke, Alexie Papanicolaou, Stephen Micinski, John Westbrook, Fred Gould

## Abstract

Adaptation to human-induced environmental change has the potential to profoundly influence the genomic architecture of affected species. This is particularly true in agricultural ecosystems, where anthropogenic selection pressure is strong. *Heliothis virescens* primarily feeds on cotton in its larval stages and US populations have been declining since the widespread planting of transgenic cotton, which endogenously expresses proteins derived from *Bacillus thuringiensis* (Bt). No physiological adaptation to Bt toxin has been found in the field, so adaptation in this altered environment could involve: 1) shifts in host plant selection mechanisms to avoid cotton, 2) changes in detoxification mechanisms required for cotton-feeding versus feeding on other hosts, or 3) loss of resistance to previously used management practices including insecticides. Here we begin to address whether such changes occurred in *H. virescens* populations between 1997-2012, as Bt cotton cultivation spread through the agricultural landscape. For our study, we produced an *H. virescens* genome assembly and used this in concert with a ddRAD-seq enabled genome scan to identify loci with significant allele frequency changes over the 15 year period. Genetic changes at a previously described *H. virescens* insecticide target of selection were detectable in our genome scan, and increased our confidence in this methodology. Additional loci were also detected as being under selection, and we quantified the selection strength required to elicit observed allele frequency changes at each locus. Potential contributions of genes near loci under selection to adaptive phenotypes in the *H. virescens* cotton system are discussed.

## Introduction

Human-induced change in the natural landscape places strong selective pressure on populations to adapt over relatively short evolutionary timescales (Palumbi 2001). These changes shape the genomes of local species, providing insight into contemporary evolution and it's implications for affected species. Cultivation of the natural landscape for agricultural purposes is one of the most ubiquitous examples of human-induced environmental change. Modern agricultural practices often involve sweeping changes to the composition of plant species across broad geographic regions, re-sculpting of the physical terrain and chemical inputs into the environment (Tilman *et al.* 2001). The strong selective pressure placed on species that inhabit agricultural ecosystems make them ideal for examining genetic responses to anthropogenic forces (Taylor *et al.* 1995).

One such major change in recent agricultural history is the commercialization of transgenic crops that themselves produce proteins for the management of key insect species. The tobacco budworm, *Heliothis virescens*, feeds primarily on cotton in its larval stages and populations in the Southern United States have been declining since the widespread planting of transgenic cotton (Supplementary Figure 1). These cotton cultivars endogenously express insecticidal proteins derived from the bacterium *Bacillus thuringiensis* (Bt), which are lethal to *H. virescens*. In the Southern United States, Bt-expressing cotton was rapidly adopted after it became commercially available for management of *H. virescens* in 1996 (James 2015; Supplementary Figure 2). Prior to the widespread use of Bt-expressing cotton, populations of *H. virescens* had evolved resistance to every insecticide used for their management (Blanco 2012), including pyrethroid insectides (Luttrell *et al.* 1987, Campanhola and Plapp 1989). Concerns over the possibility that *H. virescens* and other insect targets of Bt crops would evolve resistance to the endogenously expressed proteins spawned an entire field of research related to Bt resistance and associated genetic mechanisms (Reviewed in Heckel *et al.* 2007, Tabashnik *et al.* 2013). Of primary concern was the loss of efficacy of toxic Bt proteins (USEPA 1998, 2001, 2006). In the case of *H. virescens*, no physiological adaptation to the Bt toxin in cotton fields has been detected (Tabashnik *et al.* 2013). Yet widespread adoption of Bt-expressing crops likely placed selective pressure on *H. virescens* in other ways.

As one example, widespread planting of Bt cotton cultivars led to an overall decline in insecticide use on cotton in the United States (NASEM 2016, Benbrook 2012), including the use of pyrethroids (personal comm. D. Reisig). In 1999, North Carolina extension entomologists stopped recommending pyrethroids for the damaging generations of *H. virescens* in cotton (Bacheler 1999) and still do not recommend pyrethroids. Similarly, in Louisiana, pyrethroids are no longer recommended for *H. virescens* (LSU Ag Center 2016). In the state of California, one of the only states that makes records of pesticide sales and applications publicly available, the pounds per acre of cotton for three commonly used pyrethroids (deltamethrin, cypermethrin and cyfluthrin) declined from 0.05 to 0.03 between the years 2000 and 2012 (http://calpip.cdpr.ca.gov), and it is not clear if any of these pyrethroids sprays are currently used to target *H. virescens*. Prior to Bt cotton adoption, when pyrethroids were heavily used to suppress *H. virescens* populations, the voltage-gated sodium channel gene (*Vgsc*) was identified as one gene target of selection, and resistance-conferring alleles rose to high frequency (Park and Taylor 1997, Park *et al.* 1997). Yet in *H. virescens* and other insect species, *Vgsc* mutations often result in an overall loss of fitness for individuals carrying them (Zhao *et al.* 2000, Foster *et al.* 2005, Kliot and Ghanim 2012, Brito *et al.* 2013). Under these conditions, stability in the frequency of insecticide resistance alleles depends upon whether or not populations are continually exposed to insecticidal pressure. Therefore, one possible effect of Bt adoption in *H. virescens* is a reversion to susceptibility at their pyrethroid resistance locus.

Additional inadvertent targets of selection by Bt-expressing cotton could include loci involved in feeding and oviposition behaviors. *H. virescens* are well known for the damage they cause to cultivated cotton (reviewed in Blanco 2012), but their host plant range includes tobacco, soybean, garbanzo bean (Fitt 1989), and a number of wild hosts (Sudbrink and Grant 1995). Heritable, intra-specific variation in host choice has been observed for *H. virescens* (Sheck and Gould 1993, Sheck and Gould 1995, Karpinski *et al.* 2014), as well as other closely-related Lepidopteran species (Jallow and Zalucki 1996, Jallow *et al.* 2004, Oppenheim *et al.* 2012). It is possible that widespread adoption of Bt-expressing cotton has made cultivated cotton host plants highly toxic to and, in essence, unavailable for *H. virescens* host use. Such a scenario would select against individuals that preferentially oviposit and feed upon cultivated cotton, in favor of those that utilize alternative host plants. Allele frequency changes in genes associated with chemosensation, central nervous system function, and metabolism may have occurred as *H. virescens* was driven off of its primary cotton host plant (Blanco 2012).

In recent years, identifying genomic change in response to selective forces has been enabled by the development of next-generation sequencing (NGS) technologies. A variety of NGS-enabled marker development techniques are used to generate novel, high density marker sets for model and non-model organisms, including Restriction-site Associated DNA sequencing (RAD-seq; Baird *et al.* 2008), Genotyping-by-Sequencing (GBS; Elshire *et al.* 2011), double-digest RAD-seq (ddRAD-seq; Peterson *et al.* 2012) and others (reviewed in Andrews *et al.* 2016). These marker sets enable scientists to scan the genomes of field-collected organisms in search of the gene targets of selection, particularly where selection is strong and a reference genome assembly for read alignment is available (Lowry *et al.* 2017, Catchen *et al.* 2017). Strong selection for advantageous alleles at target genes can influence allelic composition at physically linked neutral sequences, including nearby marker sites, resulting in a genomic footprint of selection that is much broader than the target gene alone (Nielsen 2005). The breadth of this genomic footprint is influenced by several factors, including the strength of selection, the initial frequency of the advantageous allele, effective pest population size and recombination rate (Charlesworth and Charlesworth 2010).

Here we scanned the genomes of two *H. virescens* field populations collected in the Southern United States between the years 1997 and 2012 to detect loci that have changed over time. Given that pesticides and transgenic crops impose very strong selection on their target pest species (Onstad 2014), we initially focused on a genomic region known to be associated with insecticide resistance as a confirmation that ddRAD-seq could be used identify genes responsible for adaptive phenotypes under strong selection. To achieve this goal, we produced an annotated draft assembly of the *H. virescens* genome and used it for alignment of ddRAD-seq reads from barcoded individuals collected across space and time. We then tested the hypothesis that changes in a candidate pyrethroid resistance gene, *Vgsc*, could be detected through our ddRAD-enabled genome scanning techniques. Furthermore, we identified additional ddRAD-seq loci with strongly diverging marker allele frequencies, and quantified the strength of selection required to produce the observed changes at these sites. Some of the ddRAD-seq loci identified as under seleciton were linked to genes involved in toxin metabolism and chemosensation. We concluded with a discussion of the adaptive phenotypes that these newly identified gene targets of selection might produce in a field environment.

## Methods

### Insect Material

For all population genomic analyses, adult male moths were collected by pheromone-baited trap from Bossier Parish, LA, and Burleson County, TX. Collections took place in LA from May through September, and in TX from May through October, in the years 1997, 2002, 2007, and 2012. GPS coordinates from trapping locations can be found in Supplementary Table 1. Moths from each collection date and location were immediately placed together in bottles of 95% ethanol for long-term storage. Bottles from 2002, 2007 and 2012 were always held at -20°C until specimens were used, while those from 1997 were initially held at room temperature and then transferred to -20°C. To develop our *H. virescens* genome assembly, individuals from a long-standing colony strain (Gould *et al.* 1995) were sib-mated for 10 generations to produce inbred material for sequencing (Fritz *et al.* 2016). Siblings from a single inbred family were used for sequencing and analysis. Five sibling pupae were stored at -80°C prior to DNA isolation and library preparation. For all insect samples, DNA was isolated with a Qiagen Blood and Tissue Kit (Qiagen, Inc., Valencia, CA, U.S.A.) using the mouse tail protocol.

### H. virescens *Candidate Gene Approach*

A polymerase chain reaction (PCR) based upon the methods of Park and Taylor (1997) was used to amplify a 432 bp region in the alpha subunit of the *Vgsc*. The primer pair Nhp3304+ (5' ATGTG GGACT GIATG TTGGT) and Nhp3448-(5' CTGTT GAAGG CCTCT GCTAT) flanked a mutation known as L1029H. In this targeted region of the *Vgsc*, a single nucleotide polymorphism (SNP) caused a Leucine to Histidine amino acid substitution and thereby pyrethroid resistance. Additional mutations associated with pyrethroid resistance have been detected in *H. virescens*, including D1561V + E1565G and V421M (Rinkevich *et al.* 2013). We specifically targeted L1029H for our research because the D1561V+ E1565G mutations have not yet been functionally confirmed using ectopic expression assays (Rinkevich *et al.* 2013), and the V421M mutation was rarely found in our study populations, even in 1997 when phenotypic pyrethroid resistance was at its peak.

Amplicons from PCRs targeting the L1029H mutation were digested by restriction enzyme Nla-III, which cut in the presence of the resistance allele (Supplementary Figure 3). Genotypes were scored by visualizing the digested PCR products on a 3.5% agarose gel (90 to 120 min at 120 V). We examined the genotypes at this pyrethroid resistance locus for *H. virescens* individuals collected from 1997 (n = 194), 2002 (n = 204), 2007 (n = 268), and 2012 (n = 194) in LA, and 1997 (n = 142), 2007 (n = 120), and 2012 (n = 196) in TX. Changes in pyrethroid resistance allele frequencies over time and space were examined using a series of nested generalized linear regression models with binomial error structures in R version 3.1.2 (R Core Team 2014; used here and throughout). The following full model was used to examine the frequency of individual pyrethroid resistance alleles (*i*):

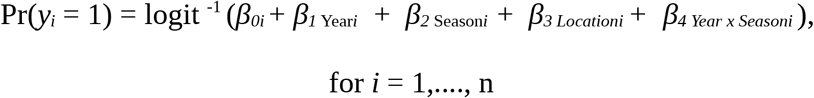
where Year represented collection year (e.g. 1997, 2002, 2007, or 2012), Season represented whether the collections were made early (May or June) or late (August through October) in the cotton growing season, and Location represented the collection location of the samples. We identified a model term as statistically significant (α = 0.05) when a comparison of nested models by analysis of deviance indicated that removal of that term significantly influenced model deviance.

### *Strength of Selection against the* Vgsc *Resistance Allele*

Following the discovery of a decline in frequency of the pyrethroid resistance allele, we quantified the strength of selection associated with the decline in pyrethroid pressure. We used the following equation to calculate the selection coefficient against the recessive resistance allele for our field populations from TX and LA over 15 years after the introduction of Bt cotton into the landscape:

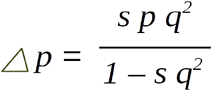
where p was the frequency of the susceptible allele, q was the frequency of the resistance allele, and *s* was the selection coefficient. To calculate the change in p over 1 generation, we took the difference in allele frequency over the 15 year period, and divided it by the total number of generations. For this, we assumed 4 generations per year (Barber 1937) for each of the 15 years examined.

### Illumina WGS Library Preparation and Sequencing

Genomic DNA from one pupa was submitted to the North Carolina State Genomic Sciences Laboratory (Raleigh, NC, USA) for Illumina paired-end (PE) library construction and sequencing. Prior to library preparation, the DNA template was quantified by a Qubit 2.0 Fluorometer (Invitrogen, USA). The PE library with an 800bp insert size was constructed using an Illumina TruSeq Nano Library Kit (Illumina, Inc. San Diego, CA) according to standard protocol. Following enrichment by PCR, the library was checked for quality and final concentration using an Agilent 2100 Bioanalyzer (Agilent Technologies, USA) with a High Sensitivity DNA chip before sequencing on an Illumina HiSeq 2500 (100×2 paired end, rapid run).

Genomic DNA from a second pupa was used for mate-pair (MP) sequencing. Prior to library preparation, whole genomic DNA was run out on a 0.5% agarose gel at 130v for 2 hours. Fragments 8kb or larger, as compared with Hyperladder I (Bioline USA Inc. Tauton, MA, U.S.A), were excised from the gel and purified using a Zymoclean large fragment recovery kit (Zymo Research Corp. Irvine, CA, U.S.A.). The DNA sample was submitted to the Michigan State University Research and Technology Support Facility (East Lansing, MI, USA) for 8kb MP library preparation and sequencing. The DNA library was prepared using an Illumina Nextera Mate Pair Sample Preparation Kit according to standard protocol. The library was validated using a Qubit dsDNA assay, Caliper LabChipGX (Perkin Elmer, Waltham, MA, U.S.A.) and Kapa Library Quantification qPCR for Illumina Libraries. The library was loaded on one lane of an Illumina HiSeq 2500 High Output flow cell and sequenced in a 2×125bp paired-end format using HiSeq SBS version 4 reagents. For both PE and MP libraries, base calling was done by Illumina Real Time Analysis v1.18.64, the output of which was converted to FastQ format with Illumina Bcl2fastq v1.8.4.

### PacBio Library Preparation and Sequencing

Genomic DNA from 4 pupae, one of which was also used for Illumina PE sequencing, were prepared into two libraries for PacBio sequencing. For each library, the SMRTbell Template Preparation Kit version 1.0 (Pacific Biosciences, Menlo Park, CA, U.S.A.) was used for gDNA preparation, but shearing and size-selection steps differed. For the first library, shearing was minimal and no size selection was performed. For the second library, shearing prior to DNA concentration was avoided to maximize fragment length, and a BluePippin (Sage Science Inc., Beverly, MA, U.S.A.) was used to select fragments that were at least 7kb long. This produced sufficient prepared library material for 17 and 5 SMRT cells, respectively. Prior to sequencing, the library concentration and fragment length profiles were checked on a Qubit 2.0 and an Agilent Tapestation 2200 (Agilent Technologies, USA) with a high molecular weight tape. Both libraries were sequenced at the University of North Carolina Sequencing facility (Chapel Hill, NC, USA) on a PacBio RS II.

### H. virescens *Genome Assembly*

Read quality was checked for all Illumina data using FastQC (Babraham Bioinformatics, Cambridge, UK). Low quality ends were trimmed from both PE and MP reads using trimmomatic (v. 0.32; Bolger *et al.* 2014) and cutadapt (v. 1.9.1; Martin 2011), respectively. Any remaining Illumina adapter sequences and Nextera transposon sequences were also removed. Reads were filtered for potential microbial contaminants and *H. virescens* mitochondrial DNA (Supplementary Data File 1) using BBmap (version 35.10; Bushnell B. - sourceforge.net/projects/bbmap/). For the full list of the screened contaminants, see Supplementary Table 2. SOAPdenovo2 (v. 2.04) was used for assembly, scaffolding and gap closure (Luo *et al.* 2012) with a k-mer length set to 63. Contigs and scaffolds over 2kb were used for further analysis.

RepeatScout (version 1.0.5; Price *et al.* 2005) was used to find *de novo*, species-specific repeats in the K63 assembly, while RepeatMasker (version open-4.0; Smit *et al.* 2013) was used to identify other common insect repeats available from Repbase (version 20150807; Jurka *et al.* 2005). We soft-masked both repeat classes using BEDTools (version 2.25.0; Quinlan and Hall 2010) and collapsed redundant haplotypes using the default settings in Haplomerger2 (version 3.1; Huang *et al.* 2012). To fill intra-scaffold gaps, we applied PacBio reads over 5kb in length to our Illumina assembly using PBsuite (version 14.9.9; English *et al.* 2012).

As an assembly quality check, BlastStation (TM Software, Inc., Arcadia, CA, U.S.A.) was used to align 654 mapped ddRAD-seq marker sequences from the F1 parent used to produce an *H. virescens* linkage map (Fritz *et al.* 2016; Dryad digital repository http://dx.doi.org/10.5061/dryad.567v8) to our scaffolds. All top hits were exported and markers with alignment hit lengths greater than 150bp (of 350 bp total), identities greater than 80%, and e-values below 0.001 were further examined. This enabled us to check for potential misassemblies, and provide additional information about which short scaffolds likely belong together on individual chromosomes (Supplementary Table 3). BlastStation was also used to identify the scaffold to which the alpha-subunit of the *Vgsc* (GenBank Accession: AH006308.2) aligned.

### Structural annotation

The Just_Annotate_My_Genome (JAMg; https://github.com/genomecuration/JAMg) platform was used to generate putative gene models. First, the genome was masked using RepeatMasker (Smit *et al.* 2013) and RepeatModeler (Smit *et al.* 2013). Subsequently, RNA-Seq data was obtained from NCBI for *H. subflexa* and *H. virescens* (SRA accessions: ERR738599, ERR738600, ERR738601, ERR738602, ERR738603, ERR738604, ERR738605, SRR1021613), preprocessed using

“justpreprocessmyreads” (http://justpreprocessmyreads.sourceforge.net), and assembled with Trinity RNA-Seq 2.1.1 (Haas *et al.* 2013) using both the ‘*de-novo’* and ‘genome-guided’ options as implemented in JAMg. The platform made use of multiple lines of evidence to support each gene model: the two Trinity RNA-Seq assemblies integrated with 63,504 publicly available Sanger-sequenced Expressed Sequence Tags using our new version of PASA (Haas *et al.* 2003); protein domain annotation of putative exons via HHblits (Remmert *et al.* 2012); the *de-novo* gene predictors GeneMark.HMM-ET (Lomsadze *et al.* 2014) and Augustus (Stanke *et al.* 2006) using the assembled and raw RNA-seq and protein domain data as external evidence. These evidence tracks were condensed to an Official Gene Set (OGS) using Evidence Modeler (Haas *et al.* 2008). A quantitative assessment of our assembly and annotation completeness was conducted using BUSCO software using the metazoan lineage setting (version 2.0.1; Simao *et al.* 2015).

### H. virescens *ddRAD-seq library preparation*

DdRAD-seq libraries were prepared according to Fritz *et al.* (2016) with minor modifications. Briefly, 200 ng of genomic DNA from the thorax of each field-collected specimen was digested with EcoRI-HF and MspI. Overhang sites from each specimen were ligated to Truseq Universal adapters (Illumina, Inc. San Diego, CA) modified to contain a unique barcode (Elshire *et al.* 2011, Fritz *et al.* 2016). Adapter-ligated DNA fragments from each individual were combined into pools of no more than 24 individuals. A Blue Pippin (Sage Science, Inc., Beverly, MA) was used to select adapter-ligated DNA fragments ranging from 450-650 bp from each pool, and size-selected DNA pools were amplified in a Peltier PTC200 thermalcycler under the following reaction conditions: 72 °C for 5min, 18 cycles of 98 °C for 30 sec, 65 °C for 20 sec, 72 °C for 30 sec, followed by 72 °C for 5 min. For each pool, 1 of 4 Illumina indices (1,2,6, or 12) was added via PCR to the MspI adapter. Amplified pools were combined, cleaned with a Qiaquick PCR Purification Kit (Qiagen, Inc., Valencia, CA, U.S.A.), and diluted to 4nM prior to sequencing. Prepared genomic DNA libraries constructed from a total of 177 *H. virescens* individuals were spread across four 2×300 paired-end Illumina MiSeq runs. Individuals from each year and collection location were spread evenly across each MiSeq run to minimize sequencing run bias in our downstream analysis.

### Demultiplexing and Genome Alignment of DdRAD-seq Markers

Illumina-generated read 1 and 2 files were merged using FLASH version 1.2.7 (Magoc and Salzburg 2011), then demultiplexed and filtered for quality using the process_radtags script from Stacks version 1.09 (Catchen *et al.* 2011, 2013). Quality filtering entailed removal of reads when: 1) they did not have an intact EcoRI cut site, 2) had a quality score < 30, or 3) were smaller than 350 bp. We disabled the rescue reads feature in the process_radtag script, and therefore no read containing errors in the barcode sequence was used for downstream analysis. All remaining merged reads were truncated to a maximum length of 350 bp. Filtered demultiplexed reads were aligned to our *H. virescens* genome assembly using Bowtie 2 (version 2.2.4; Langmead and Salzberg 2012). All reads were aligned in end-to-end mode using the preset parameters with the highest sensitivity (--very-sensitive).

### Association of DdRAD-seq Marker Genotypes with the Pyrethroid Resistance Allele

We first identified whether any raw ddRAD sequencing reads aligned to the scaffold containing the *Vgsc* using Integrative Genomic Viewer (IGV; Robinson *et al.* 2011). Following identification of potential ddRAD-seq markers near the *Vgsc*, we inspected stacks of ddRAD-seq reads for individuals with genotypic data at the pyrethroid resistance locus. Particular attention was paid to individuals that were homozygous for the pyrethroid resistance allele. Through an IGV visual inspection of ddRAD-seq raw reads, we identified one 350bp ddRAD-seq locus (hereafter Hv_11322), for which a single 350bp sequence (hereafter Hv_11322_hap1) was commonly associated with the L1029H mutation at the *Vgsc*. Filtered, genome-aligned reads from all specimens were then fed into the Stacks v. 1.09 (Catchen *et al.* 2011, 2013) pipeline for read clustering. Custom R and python scripts were used to call 350bp ddRAD-seq genotypes at Hv_11322 for all field-collected individuals, which were then manually inspected and edited to include any insertions and deletions that were omitted by the Stacks software. For purposes of genotype calling at Hv_11322, individuals with a read count of 6 or higher for a single 350bp sequence were considered homozygotes, with two copies of that allele. Where individuals carried fewer than 6 reads for a single 350bp sequence, their genotypes were scored as a single copy of that observed allele plus one null allele. This threshold was chosen because individuals with 6 or more reads can be called homozygotes with greater than 95% certainty (Buerkle and Gompert 2013).

We postulated that if the breadth of the “selective sweep” surrounding the *Vgsc* resistance allele included Hv_11322, such that Hv_11322_hap1 was associated with the L1029H mutation, the rates of their decline in frequency should be similar, if indeed there was a decline. We therefore examined whether the frequencies of the L1029H mutation and Hv_11322_hap1 differed in their rate of decline over time. Specifically, we used a series of nested generalized linear models with binomial error structures to examine whether locus and collection year interacted to influence individual allele (i). In the case of the Hv_11322 response, Hv_11322_hap1 was scored as a 1 and all other alleles were scored as a zero. Our full statistical model was as follows:

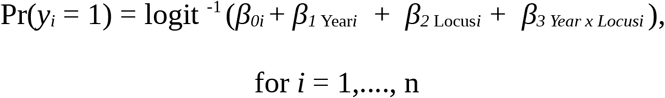
where Year represented the years during which the moths were collected and Locus indicated either the *Vgsc* or ddRAD-seq marker Hv_11322. As before, we identified a model term as statistically significant when a comparison of nested models by analysis of deviance indicated that removal of that term significantly influenced model deviance. No significant difference between a model with and without the interaction term indicated that the slope of the decline in the L1029H mutation was similar to that of Hv_11322_hap1.

We also analyzed the distribution of Hv_11322_hap1 for groups of individuals that were homozygous for either the resistant or susceptible alleles at the *Vgsc*. In total, 32 individuals were homozygous in our target region of the *Vgsc* and contained sufficient ddRAD-seq data at nearby locus Hv_11322 to call at least one allele. Of these 32 individuals, two Hv_11322 alleles could be called for 26 individuals, whereas only a single allele could be confidently called for 6 of the individuals due to their lower than 6X depth of coverage. In total, 58 haplotypes (from 32 individuals), which contained genotypic information for both the *Vgsc* and the nearby ddRAD-seq marker were examined. A Fisher's exact test of independence was used to determine whether there was an association between the frequencies of Hv_11322_hap1 and the L1029H mutation.

### H. virescens *ddRAD-seq Enabled Genome Scan*

Previous work by Groot *et al.* showed that genetic differentiation among North American populations of *H. virescens* was low (Groot *et al.* 2011). Given these already documented high levels of gene flow between collection sites, and our goal to detect genomic change over time, we specifically focused on analyzing allele frequency changes between years. Samtools (version 0.1.18; Li *et al.* 2009, Li 2011) view was used to convert SAM files output by Bowtie 2 to BAM files, and SNPs were called using mpileup. BCFtools was used to generate SNP and indel genotypes, as well as genotype likelihoods in a Variant Call Formatted (VCF) file. This VCF file was filtered by VCFtools (version 0.1.15, Danacek *et al*. 2011, https://vcftools.github.io) prior to downstream population genomic analysis. The filtered dataset included loci that: 1) were sequenced to a depth of 3 or more reads, 2) had a minor allele frequency of 0.1 or greater, 3) were represented in at least 50% of individuals, and 4) included only SNP variant sites (indels were excluded). The number of SNPs was thinned such that no more than 2 were examined per ddRAD-seq locus. This thinned SNP dataset was transformed from VCF format to genepop format using PGDSpider (version 2.1.1.0, Lischer and Excoffier 2012). Pairwise SNP outlier analyses of these thinned SNP datasets were made using Lositan (Antao *et al.* 2008) with the following parameter settings: 1) “neutral” and “forced” mean FST settings were engaged, 2) the Infinite Alleles model was assumed, 3) the false discovery rate was set to 0.1, and 4) the type I error was set to 0.01.

Scaffolds containing SNP markers with statistically significant pairwise-genetic divergence were identified for further analysis (Supplementary Data File 2). Using the physical distance between the ddRAD-seq locus Hv_11322 and the *Vgsc* as our guide, the predicted structural genes on each scaffold found within 36 kb of each SNP outlier were examined. Protein sequences corresponding to each annotation along a scaffold where divergent markers were present were aligned to the NCBI Arthropod database (taxid: 6656) via blastp using Blast2GO software.

Given the large number of outlier loci (Figure 3), and that the potential traits under selection in this agricultural system (e.g. metabolic detoxification of insecticides and/or host plant defensive compounds, host volatile detection) are often quantitative, we reasoned that certain gene families may be over-represented near our outliers with respect to their overall distribution throughout the *H. virescens* genome. Therefore, we examined the distribution of GO categorizations (54 level two categorizations in each of three GO domains: “Biological Process”, “Molecular Function”, and “Cellular Component”) for the subset of predicted genes found near outlying SNPs for each by-year comparison. We compared those distributions to the numbers of genes found in each of these same categories in the overall *H. virescens* genome using a series of Fisher's Exact tests. Due to the large number of comparisons (n = 54), we used a Bonferroni-corrected alpha value of 0.0009 to establish statistically significant over-representation in any one GO category. For each by-year comparison, we examined the subsets of genes within 2 different intervals from SNP outliers: 10 kb (“moderate” linkage according to Lowry *et al.* 2017) and 36 kb (extended linkage, based upon the distance between Hv_11322 and the *Vgsc* target of selection).

### Strength of Selection on Outlier Loci

Finally, we examined what the strength of selection must have been to produce the observed allele frequency changes at all identified outlier loci for each by-year comparison. We calculated the coefficient of selection (*s*) against q, the declining allele frequency, assuming: dominance of p, recessiveness of p, or incomplete dominance of p (where h = 0.5) according to Falconer and Mackay (1996). Custom scripts written in R were used to calculate *s* for all outliers with the exception of those with initial values of p that were very low (p > 0.05). These were excluded because they are most susceptible to under-sampling due to our small sample sizes, and small biases in these values have the potential to significantly influence the selection coefficient.

## Results

### H. virescens *Candidate Gene Analysis*

In 1997, the frequency of the L1029H mutation was 0.66 in LA and 0.63 in TX. By the year 2012, the frequency of this resistance allele declined to 0.44 in LA and 0.36 in TX (Figure 1). This decline in the resistance allele frequency over our 15 year sampling period was statistically significant (p < 0.001). Neither the interaction between year and season (p = 0.36), season itself (p = 0.21), nor sampling location (p = 0.25) significantly influenced the frequency of the resistance allele.

**Figure 1.**
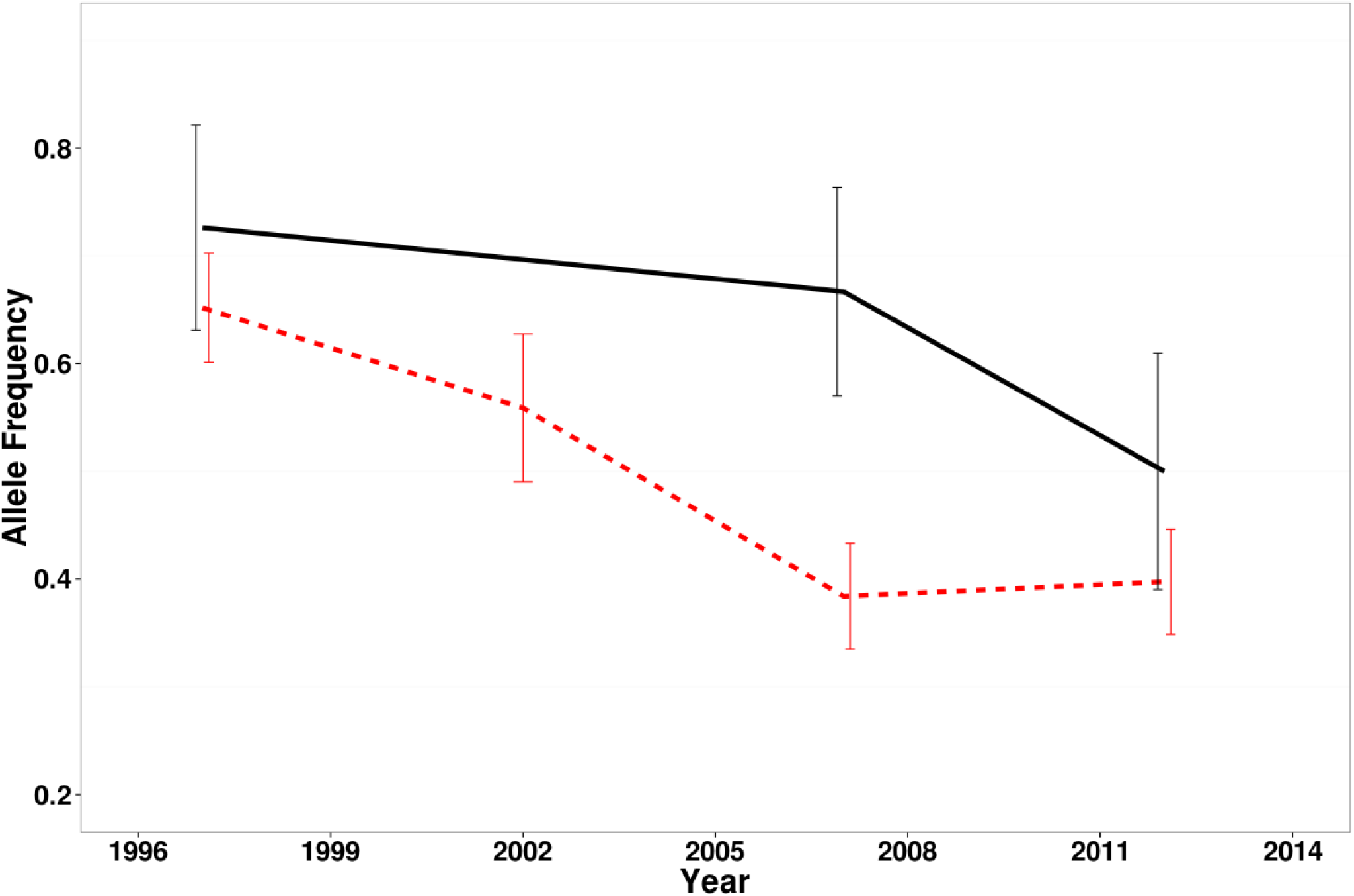
The decline in the frequency of the pyrethroid resistance allele in *H. virescens* (pooled LA and TX samples), represented by the dashed red line, was statistically significant over the course of our 15 year sampling period (n = 659). A unique ddRAD-seq haplotype, represented by the solid black line, was found *ca.* 36Kb upstream from the alpha subunit of the *Vgsc* and also declined in frequency in the subset of individuals (n = 141) sequenced for our genome scan. Error bars represent bootstrapped 95% confidence intervals (N = 5000) around the mean of each year.

### Strength of Selection against the Vgsc Resistance Allele

In LA, the frequency of the resistance allele declined by 0.22 over the 15 years between 1997 and 2012. In TX, the decline in the resistance allele frequency was 0.27. Therefore, the increase in p per generation was 0.004 and 0.005 for LA and TX, respectively. With this information, we were able to calculate a selection coefficient of 0.03 for each of the LA and TX populations.

### Genome Sequencing and Assembly

In total, the Illumina sequencing runs produced 122,433,923 and 232,607,659 reads for PE and MP libraries, respectively. After read trimming and filtering, 115,374,414 and 227,857,423 reads from the PE and MP libraries were used for assembly. An additional 482,464 PacBio reads with an average length of 7560 (s.d. = 2663) bp were applied to our Illumina assembly using PBsuite software for gap filling. The final *H. virescens* genome assembly was comprised of 8,826 scaffolds with a total length of 403,154,421 bp, similar to the previously estimated *H. virescens* genome size of 401 Mbp (Gregory and Herbert 2003). The scaffold N50 was 102,214 bp (mean size = 45,678 bp; range = 659 – 628,964 bp). A BUSCO analysis of our final assembly indicated that 865 (88%) of the 978 core conserved eukaryotic genes were complete. Further specifications for our genome assembly can be found in Supplementary Table 4. When we examined our previously mapped *H. virescens* ddRAD-seq markers (Fritz *et al.* 2016), a total of 562 out of 654 met the aforementioned alignment criteria relative to the reference genome and were used to examine and group scaffolds into chromosomes. Of these 562 markers, 557 aligned uniquely to a single scaffold, while 5 markers (4851, 5891, 13906, 22644, 29612) aligned well to multiple scaffolds. This suggested that either those scaffolds were allelic, or that the marker sequences contain repetitive DNA. Four-hundred eighty three of the 8826 scaffolds present in our assembly were aligned to at least 1 mapped marker. In most cases (n = 421 scaffolds), a single scaffold was associated with a single mapped marker. However, 62 scaffolds could be aligned to multiple mapped markers, which enabled us to check the quality of our assembly against our linkage map. Of these 62 scaffolds, only 5% (3 scaffolds) aligned to markers that originally mapped to different linkage groups. A summary of the scaffold names and groupings by linkage group can be found in Supplementary Table 3. One scaffold, numbered 4600, contained the entire *Vgsc* sequence available from GenBank accession AH006308.2.

### Association of DdRAD-seq Marker Genotypes with the Pyrethroid Resistance Allele

We located one ddRAD-seq marker, called Hv_11322, spanning bp 11,397 through 11,747 of Scaffold 4600 which is *ca.* 36 kb upstream from the pyrethroid resistance locus. In total, 55 unique Hv_11322 sequences (alleles) could be identified from 138 individuals using Stacks. Fifty of these 55 alleles were found fewer than three times in our field-collected populations. The remaining five most common alleles (Supplementary Figure 4) were found 5, 5, 6, 12, and 161 times, respectively. According to an NCBI blast, all of these sequences aligned well with GenBank accession DQ458470.1, a DNA sequence from *Helicoverpa armigera* that contained the *Vgsc*.

No statistically significant difference existed between the slope of the decline in the L1029H mutation and Hv_11322_hap1 (deviance = -0.355, df = 1, p = 0.551). These allele frequency declines are plotted in Figure 1. When we examined the full haplotypes (e.g. containing both the Hv_11322 locus and the pyrethroid resisance locus), specifically in homozygotes at the pyrethroid resistance locus, we identified 20 unique Hv_11322 alleles in the 32 total individuals (or N = 58 chromosomes). When chromosomes containing the L1029H mutation (pyrethroid resistance-conferring *Vgsc* allele) were examined, 85% (29 of 34) also carried the ddRAD-seq allele Hv_11322_hap1 (Figure 2). The remaining 5 chromosomes bearing the L1029H mutation contained 4 unique Hv_11322 alleles, none of which were the 5 most common Hv_11322 alleles. Only 5% (1 of 24) of the chromosomes bearing the wild-type *Vgsc* allele also carried Hv_11322_hap1. Fifteen unique Hv_11322 alleles of the 20 alleles found in homozygotes were associated with the wild-type *Vgsc* allele. A Fisher's Exact test indicated that there was a statistically significant association between the presence of Hv_11322_hap1 and the L1029H mutation (p < 0.001).

**Figure 2.**
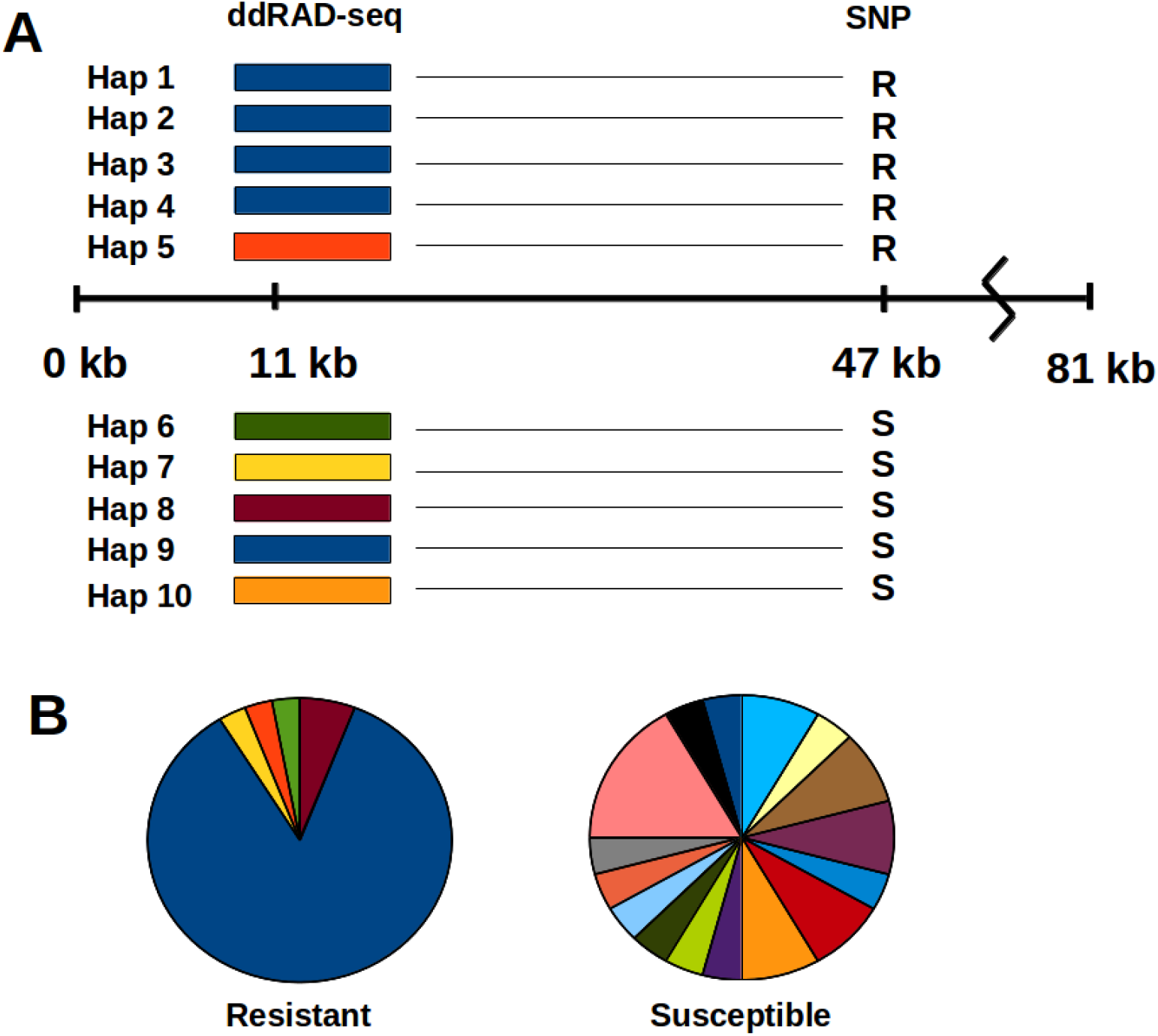
Significantly greater diversity was observed from ddRAD-seq haplotypes linked to the susceptible *Vgsc* allele relative to those linked to the resistance allele. (A) is a visualization of the relationship of Hv_11322 to the L1029H *Vgsc* SNP along an 81kb genome scaffold. The different colored bars at 11kb represent unique alleles at the Hv_11322 locus, and depict the greater diversity of Hv_11322 alleles associated with the *Vgsc* wild-type allele, relative to the resistance allele. Due to the number of unique Hv_11322 alleles associated with each *Vgsc* allele (n = 5 and n = 15 for resistant and wild-type, respectively), full representation of this diversity could not be incorporated into (A). However, the unique colors in (B) depict the number and proportion of Hv_11322 alleles linked to the *Vgsc* resistant (n = 34) and susceptible (n =24) SNP alleles. Each unique color represents a different Hv_11322 allele. The dark blue wedges always represent Hv_11322_hap 1.

**Figure 3.**
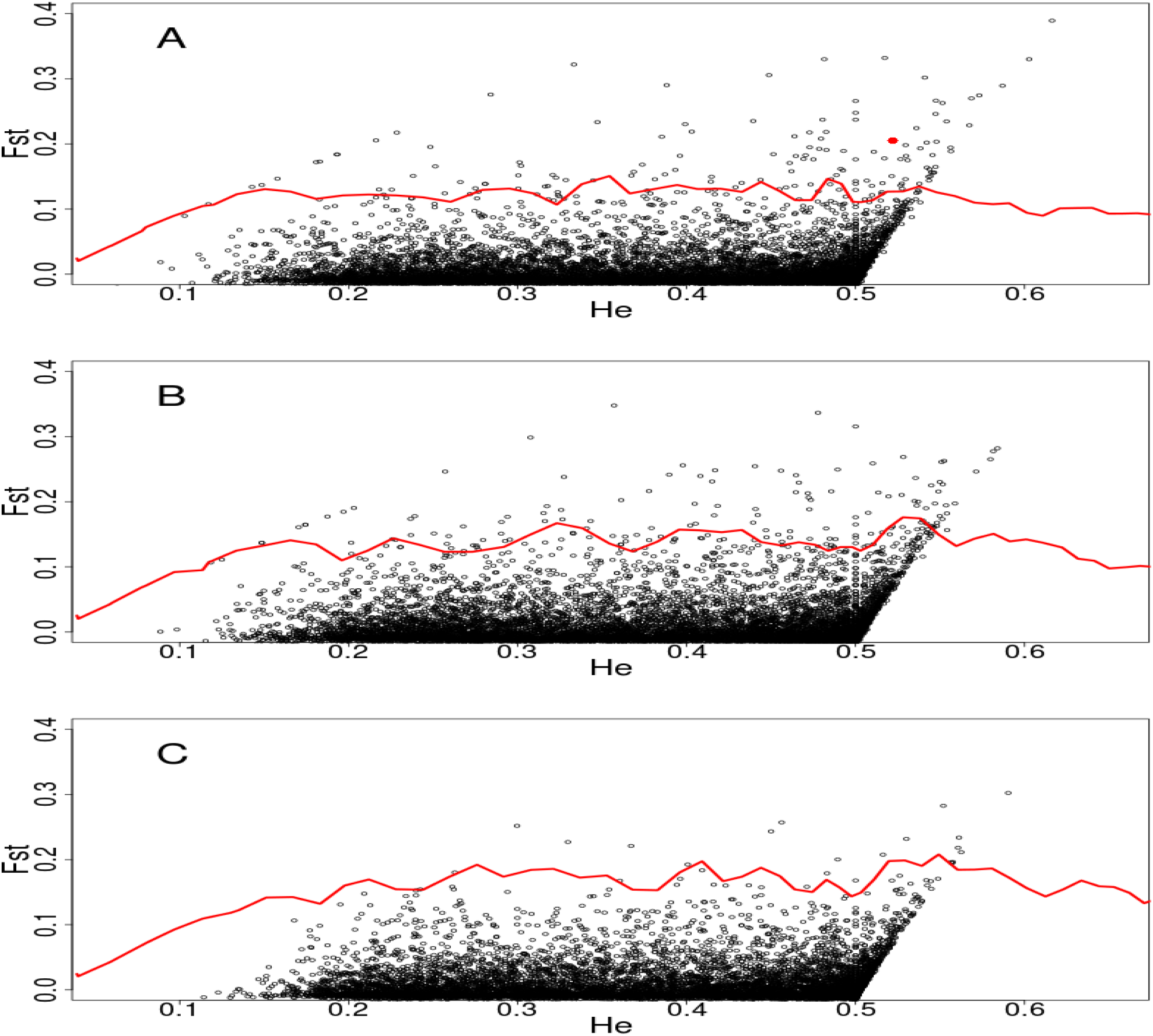
Pairwise genetic divergence according to Weir and Cockerham's FST for populations of *H. virescens* collected in the years A) 1997 and 2012, B) 1997 and 2007, C) 2007 and 2012. Each black point represents one SNP of 8,963 along the *H. virescens* genome. Points above the red line represent loci with pairwise genetic divergence that is statistically significant at the α = 0.01 level following correction for false discovery. Pairwise genetic divergence at the SNP near the *Vgsc* on Scaffold 4600 is represented by the red point on panel A.

### H. virescens *ddRAD-seq Enabled Genome Scan*

Of the 1,682,114 SNPs in our ddRAD-seq marker dataset, the total number of filtered SNPs included in the analyzed dataset was 8,963. Based upon this filtered dataset, overall population genomic divergence between years was low. Pairwise Weir and Cockerham's overall FST values were 0.005, 0.004, and 0.013 for the comparisons 1997-2007, 1997-2012, and 2007-2012, respectively.

We first examined SNPs along Scaffold 4600, where the *Vgsc* was located, for evidence of genomic divergence between years. Between the years 1997 and 2012, one SNP at position 11706 on Scaffold 4600, which was part of Hv_11322, showed signs of statistically significant genetic divergence with a Weir and Cockerham's FST value of 0.205 (p < 0.001). This SNP, a cytosine to thymine transition, went from a cytosine allele frequency of 0.875 (n = 44 diploid individuals) in 1997, fell to a frequency of 0.724 in 2007 (n = 49) and further declined to a frequency of 0.548 in 2012 (n = 52; Fig. 1).

In addition to this FST outlier on Scaffold 4600, our genome scan revealed a number of additional diverging allele frequencies for each by-year comparison. In total, we detected a total of 351 SNPs on 314 scaffolds (3.6% of the 8,826 total scaffolds) as outliers in at least one by-year comparison. Table 1 shows the number of genomic outliers for each comparison, as well as the number of unique scaffolds on which these unique outliers were found. Between the years 1997 and 2007, 201 SNPs (2.2% of the 8,963 total SNPs examined) showed signs of significant allele frequency divergence, whereas only 35 SNPs (0.4% of the 8,963 total SNPs) significantly diverged between the years 2007 and 2012. When a comparison was made over the total time period, between the years 1997 and 2012, 184 SNPs showed signs of statistically significant allelic divergence.

**Table 1.**
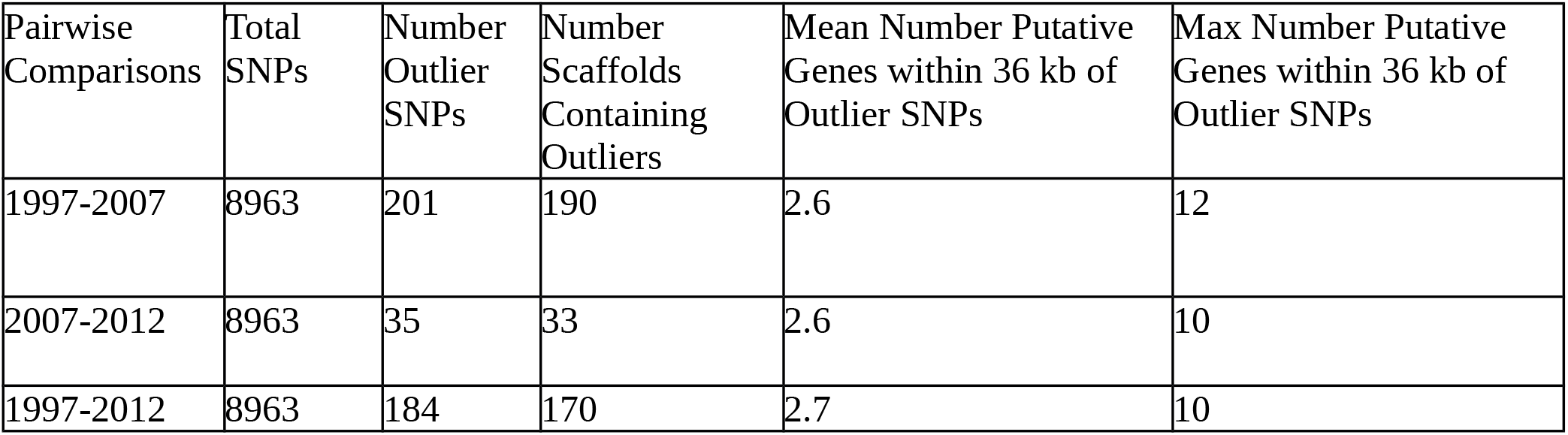
Number of SNPs with pairwise FST values deemed as significantly divergent according to Lositan analysis for each by-year comparison. The numbers of unique scaffolds (of 8826) containing at least 1 significantly diverged SNP, as well as the average and maximum numbers of putative genes within 36 kb of the outlier SNP are also included.

| Pairwise Comparisons | Total SNPs | Number Outlier SNPs | Number Scaffolds Containing Outliers | Mean Number Putative Genes within 36 kb of Outlier SNPs | Max Number Putative Genes within 36 kb of Outlier SNPs |
| --- | --- | --- | --- | --- | --- |
| 1997-2007 | 8963 | 201 | 190 | 2.6 | 12 |
| 2007-2012 | 8963 | 35 | 33 | 2.6 | 10 |
| 1997-2012 | 8963 | 184 | 170 | 2.7 | 10 |

In some cases, the same SNPs showed signs of divergence in two different by-year comparisons, but no SNP outliers were shared among all 3 comparisons (Supplementary Figure 5). Fifty-three SNPs were considered outliers between the years 1997 and 2007, and again between 1997 and 2012. This indicated that significant allele frequency changes occurred between the years 1997 and 2007, followed by stability or small, non-significant allele frequency changes through the year 2012. Likewise, 5 SNPs showed significant allele frequency divergence between the years 1997 and 2012, and again between the years 2007 and 2012, indicating that allele frequencies were stable, or underwent modest changes between the years 1997 and 2007 followed by a significant change in the year 2012. A complete list of SNP outliers and their genome positions for each by-year comparison can be found in Supplementary Data file 2.

When we examined the DNA flanking the SNP outliers (up to 36 kb on either side for a total of 72 kb), the mean number of putative genes identified in these broad genomic regions was fewer than 3, and the maximum was 12 (see Table 1 for a complete breakdown by comparison). Some outliers were found near putative genes with functions related to either insecticide resistance or changes in host use. For example, one predicted gene sequence within 10kb of two outliers, JAMg_model_7840, aligned to a cytochrome p450 protein sequence (*Cyp6AE12*) from *Helicoverpa armigera* (GenBank Accession AID54888.1) with a 100% query cover and 83% identity. It was identified on Scaffold 3424, and the associated SNPs were detected as outliers between 1997-2012, and also 2007-2012. As a second example, two predicted gene sequences, JAMg_model_4651 and JAMg_model_4652 aligned with at least 70% query cover and 78% identity to *Athetis lepigone* olfactory receptor (OR60; GenBank Accession KT588155.1) and *Helicoverpa assulta* olfactory receptor (OR33; GenBank Accession KJ542684.1) sequences, respectively. These gene sequences were found within 10kb of a SNP outlier from 1997-2007 and 1997-2012 comparisons and was located on Scaffold 2173. We examined the ontology of genes near SNP outliers, with an eye toward those predicted to be important for detoxification and behavior, to identify whether any GO category was over-represented near outlier SNPs relative to the overall genome.

GO assignments by Blast2GO were only achieved for a subset of predicted genes. The proportions of putative genes within 36 kb of outliers for which a function could be predicted were 0.37, 0.38, and 0.32 for the by-year comparisons 1997-2007, 1997-2012, and 2007-2012, respectively. While low, these were similar to the proportion of gene predictions assigned a function by Blast2GO in the overall genome assembly (0.40). When Fisher's Exact tests were applied to identify whether any level 2 GO term was over-represented in either 10 or 36kb windows near outlier SNPs from each by-year comparison, no significant difference was found relative to their distribution in the overall genome.

### Strength of Selection on Outlier Loci

Selection coefficients (*s*) were calculated for 164, 150, and 31 outliers for the following by-year comparisons, respectively: 1997 and 2007, 1997 and 2012, and 2007 and 2012. For selection against q, the declining allele, *s* ranged from 0.009 – 0.294 across all by-year comparisons when dominance of p was assumed. When incomplete dominance of p was assumed, selection coefficients ranged from 0.012 to 0.291. Finally, when recessiveness of p was assumed, selection coefficients ranged from 0.007 – 0.737.

The mean selection coefficients (± standard deviation) calculated across outliers were 0.045 (0.018), 0.073 (0.027) and 0.147 (0.055) assuming dominance of p for the 1997 - 2012, 1997 - 2007, and 2007 - 2012 by-year comparisons, respectively. Mean selection coefficients assuming incomplete dominance of p were 0.052 (0.027), 0.074 (0.036), and 0.142 (0.060) for these same time periods. The assumption of a recessive p led to even greater mean selection coefficients; 0.129 (0.147), 0.159 (0.172), and 0.232 (0.209) during 1997-2012, 1997-2007, and 2007-2012, respectively. In general, SNPs with higher selection coefficients were associated with greater rates of change in the frequency of q, as well as higher initial starting values of q (Figure 4).

**Figure 4.**
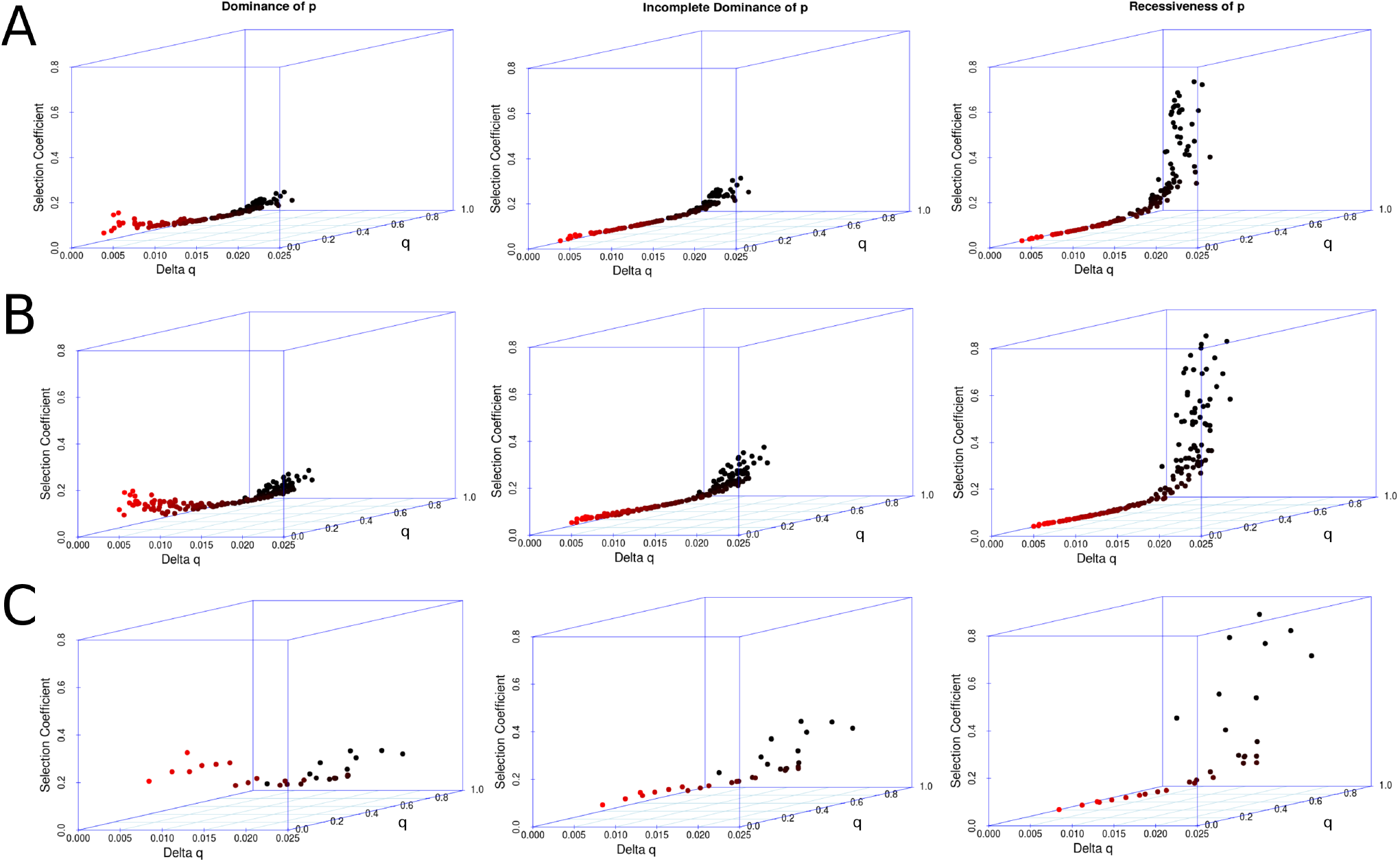
Selection coefficients associated with SNP outliers where the initial frequency of p was greater than 0.05. Selection against q is plotted against the rate of allele frequency change and the initial frequency of q. Colors for each plotted point help to visualize the initial frequency of q, where red is low and black is high. Plots in rows A, B, and C represent selection coefficients assuming different degrees of dominance of p (dominance, incomplete dominance, recessiveness from left to right) for each of the by-year comparisons 1997-2012, 1997-2007, and 2007-2012, respectively.

## Discussion

Double-digest RAD-seq and other NGS marker-development methods have been used to detect signatures of local adaptation in a number of non-model plant and animal species (e.g. Hohenlohe *et al*. 2010, Nadeau *et al*. 2014, Pujolar *et al.* 2014, Ruegg *et al.* 2014, Pais *et al.* 2016). Here we have demonstrated the power of ddRAD-seq to identify genomic regions that have diverged over short evolutionary time scales (fewer than two decades) in a landscape characterized by human-induced environmental change. We postulated that widespread adoption and cultivation of Bt cotton in the Southern United States would likely impose strong selection on Lepidopteran herbivore and cotton pest, *H. virescens*, through shifts in host plant composition and insecticide use. We first identified allele frequency changes at a likely gene target of selection, the *Vgsc*, in field-collected populations of *H. virescens.* We then demonstrated that this change could be detected using a nearby ddRAD-seq marker. Allele frequencies at many other regions of the *H. virescens* genome also diverged over time, likely in response to selection pressures imposed by widespread adoption of Bt cotton. We calculated selection coefficients for SNPs that were detected as having changed significantly over time, to demonstrate the strength of selection encountered by organisms found in agricultural ecosystems. Furthermore, we sequenced and assembled an *H. virescens* genome to help us identify potential structural genes involved in adaptation to agricultural inputs, and made it publicly available at NCBI.

Our initial examination of the *Vgsc*, a candidate gene likely to be impacted by the decline in pyrethroid use that followed Bt cotton adoption, demonstrated that the resistance-conferring L1029H mutation declined in frequency over time. Indeed, examination of allele frequency changes at the *Vgsc* locus yielded a selection coefficient of 0.03, indicating that the resistance allele was deleterious in field populations where pyrethroid pressure was low. A selection coefficient of this magnitude seemed reasonable given the previously reported fitness cost associated with carrying this resistance allele (Zhao *et al.* 2000). We were surprised to see the frequency of the resistance allele plateau in the year 2007 and remain at *ca.* 0.4 through the year 2012, however. There are several possible explanations for this. Perhaps pyrethroid use declined, but because it was not eliminated from the agricultural landscape, selection still maintains the resistance allele at lower frequency in field populations. Alternatively, the fitness cost associated with carrying the resistance allele in the absence of pyrethroid selection may only manifest itself in homozygotes. Under these conditions, the resistance allele could be maintained in heterozygotes, making it difficult to purge from *H. virescens* populations.

Using a ddRAD-seq dataset, we identified one marker that aligned to our reference genome *ca.* 36 kb upstream of the *Vgsc*. One allele of this 350bp marker, called Hv_11322_hap1, was associated with the L1029H mutation that confers pyrethroid resistance. This suggested that Hv_11322_hap1 was in linkage disequilibrium with the L1029H mutation. Furthermore, the breadth of the selective sweep in this genomic region extended at least 36 kb on one side of the *Vgsc*. Upon further examination of Scaffold 4600, which contains this region under selection, we identified three cytochrome p450s that are found between Hv_11322 and the *Vgsc*. This confirmed previous reports of tight physical linkage between the *Vgsc* and *Cyp6B10* in *H. virescens* (Park and Brown 2002). It is possible that these cytochrome p450s could also be targets of selection by pyrethroid insecticides, and future work could be directed at whether or not they play any roles in the expression of pyrethroid resistance phenotypes. Work in another closely-related Lepidopteran species, *H. armigera*, suggests that this particular cytochrome p450 is not involved in pyrethroid resistance, however (Grubor and Heckel 2007).

SNP data from our ddRAD-seq marker Hv_11322 enabled us to rediscover changes at the *Vgsc* associated with the L1029H mutation over time. While the SNP outlier in the Hv_11322 marker demonstrated significant allelic divergence relative to the genome-wide average FST value, additional SNP outliers on other *H. virescens* scaffolds were also detected. In spite of the fact that we applied a correction to reduce detection of false positives, it is possible that up to 35 of our total 351 SNP outliers are false positives given our false discovery rate of 0.1. This correction threshold was selected to minimize false positive detection, while retaining true positives (Verhoeven *et al.* 2005). To safeguard against pursuing potential false positives, further research could initially focus on genes near outliers with significant allele frequency changes in multiple by-year comparisons. Fifty-three SNPs showed significant allelic divergence across multiple by-year comparisons, where major allele frequency shifts took place between the years 1997 and 2007. Between 2007 and 2012, 5 SNPs showed significant shifts in allele frequency and these changes were detected in multiple by-year comparisons as well. It is likely that these SNP markers are in linkage disequilibrium with gene targets of selection as management of cotton ecosystems has led to the replacement of conventional cotton cultivars with Bt-expressing varieties. Interestingly, two of these SNPs were found on scaffolds containing genes with plausible roles in detoxification (of insecticides and plant defensive compounds) or host plant detection.

Scaffold 3424 contained SNPs that diverged significantly over time in our field-collected populations of *H. virescens*. Divergence was strongest in by-year comparisons from 1997-2007, and 1997-2012. This suggests that most genomic change occurred between the years 1997 and 2007, and that allele frequencies remained stable between the years 2007 and 2012. Blast results for predicted gene sequences found on this scaffold revealed homology with the cytochrome p450 superfamily. The predicted sequence aligned well with an *H. armigera Cyp6AE* gene, which is a cytochrome p450 family known to be involved in detoxification (Zhou *et al.* 2010). The best alignment was to an *H. armigera Cyp6AE12,* and expression levels of this gene in *H. armigera* are modified in response to pyrethroid insecticides (Yue *et al.* 2007, Zhou *et al.* 2010). It is possible that allelic changes on this scaffold are a response to reduced pyrethroid use in the Southern United States as a result of Bt cotton deployment in the agricultural landscape.

An alternative explanation exists for changes in allele frequencies at this candidate gene, however. *Cyp6AE12* expression is also modified in response to the plant compound xanthotoxin (Zhou *et al.* 2010). Another *Cyp6AE* monoxygenase, *Cyp6AE14,* has 68% protein sequence similarity to *H. armigera Cyp6AE12*, is highly expressed in the midgut of *H. armigera,* and is likely involved in detoxification of the cotton defensive compound gossypol (Mao *et al.* 2007). Indeed, RNAi silencing of *Cyp6AE14* in *H. armigera* led to a decline in larval growth when gossypol was present in their diet (Mao *et al.* 2007). If this *H. virescens* cytochrome p450 is indeed the target of selection, it is possible that the divergence in allele frequencies between the years 2007 and 2012 was the result of selection for improved larval performance in *H. virescens* populations that feed on alternative hosts. When widespread planting of Bt-expressing cotton drove *H. virescens* off of their previously abundant cotton host to alternative host plants, *H. virescens* would have been exposed to new plant defensive compounds. Genes associated with larval performance in one host plant genera may or may not be associated with improved larval performance on other host plant genera (Sheck and Gould 1993). Therefore, allelic changes near our *H. virescens* SNP outlier may be caused by relaxation of selection for gossypol detoxification as they moved out of cotton and adapted to new host plants and their antifeedant chemicals. To determine whether phenotypes resulting from these molecular shifts are directly associated with changing *H. virescens* management practices (e.g. pyrethoid or Bt toxin use) or shifts in host plant use, further work could involve measuring associations between genotypes at target genes on these scaffolds and the aforementioned phenotypes of interest.

Equally as important to host shifts is the ability of adult females to find a suitable host for oviposition. Chemosensation is important for host plant identification in phytophagous insects, and genes involved in olfaction and gustation are often targets of selection when host shifts occur (Dworkin and Jones 2009, Smajda *et al.* 2012). An outlier SNP was detected on *H. virescens* Scaffold 2173 that underwent allele frequency changes between the years 1997 and 2007, then remained stable between the years 2007-2012. Within 10 kb of this outlier was a pair of olfactory receptor genes. It is plausible that allele frequency changes at these olfactory receptor genes may reflect changes in the olfactory percept, enabling females to identify non-cotton host plants. Future work directed at elucidating the phenotypic effects of changes at these genes will be critical to determining their role, if any, in host plant detection.

Both toxin metabolism and chemosensation can be complex traits with multiple loci involved. Arrays of duplicated cytochrome p450s (Li *et al.* 2002, David *et al.* 2013) and carboxylesterases (Field *et al.* 1988, Guillemaud *et al.* 1997) have been implicated in metabolic detoxification of chemicals in other insect species. Likewise, clusters of olfactory and gustatory receptors are thought to contribute to host-plant utilization in some insect species (Smajda *et al.* 2012). Therefore, we examined GO level 2 categorizations, particularly those related to detoxification and chemosensation, for the putative genes surrounding our SNP outliers to determine if any GO categories were over-represented near our outliers relative to the rest of the genome. Our results did not suggest there was any statistically significant over-representation of any GO category near our outliers for any of our by-year comparisons. There are at least two possible explanations for this: 1.) The targets of selection linked to our SNP outliers do not necessarily involve duplicated, amplified, or arrayed genes of similar function. Instead these targets may involve single copy genes or gene regulatory regions associated with adaptive phenotypes in *H. virescens*. 2.) Only 30-40% of the putative genes in our genome were assigned a function in our annotation pipeline. Over-representation of certain GO categories may exist, but because 60-70% of putative genes in our genome could not be assigned a function, we were unable to detect it.

While our *H. virescens* draft assembly was instrumental to identifying selection at the *Vgsc*, as well as identifying novel genomic regions under selection in field-collected populations, there is room for improvement. For example, some SNP outliers were found on scaffolds for which there were few putative genes. These scaffolds were often short, which prohibited detection of candidate genes linked to these SNP outliers. Future work aimed at enhancing our assembly contiguity would not only improve our ability to detect candidate genes near SNP outliers, but would also enable chromosome-level identification of runs of homozygosity, a hallmark of selection. As previously mentioned, some outliers were found near putative genes, but these putative genes could not be assigned a function. Future manual curation efforts will improve our official gene set and the likelihood of identifying additional gene targets near our SNP outliers.

While the gene targets of selection identified during our study period require further validation, the strength of selection near these targets, as measured by *s,* ranged from 0.009 and 0.737, depending upon by-year comparison and degree of dominance assumed. Following theoretical expectations, selection against q required higher selection coefficients when initial frequencies of q were high (Figure 4). Furthermore, degree of dominance impacted *s.* When incomplete dominance rather than complete dominance of p was assumed, values of *s* were, on average, 7-21% higher depending upon by-year comparison. These values increased further (by 85-200%, on average) when recessiveness, rather than dominance, of p was assumed. Indeed, examples of selection against a dominant susceptible allele (q), resulting in the increase in frequency of a recessive resistance allele (p) have been described in the insecticide resistance literature (for example, see Ferré and Van Rie 2002).

Across all outlier SNPs discovered within each by year comparison, the average selection coefficients ranged from 0.045 – 0.232. Interestingly, these average selection coefficients were similar to those calculated in other Lepidopteran study systems: *Biston betularia*, *Heliconius melpomene,* and *Heliconius erato*. Multiple approaches were used to estimate strength of selection on the distinct color morphs of each species, as imposed by avian (visual) predators. Whether assessing differential predation pressures experienced by spatially segregated populations or within populations over time, coefficients ranged from 0.1-0.23 and were described as the result of strong selective pressure (Mallet and Barton 1989, Mallet *et al.* 1990, Linnen and Hoekstra 2009). Ultimately, our similar average values of *s* underscore the strong selection imposed on species found in agricultural ecosystems.

In conclusion, we demonstrated that ddRAD-seq enabled genomic scanning can be used to identify organismal responses to anthropogenic changes in agricultural ecosystems, even on short, 15 year time scales. We identified additional genomic regions in this Lepidopteran species that are likely changing in response to shifts from conventional cotton planting to widespread Bt-cotton adoption.

From an applied perspective, our results suggest that ddRAD-seq genome scans may be useful for monitoring pest populations for real-time changes in allele frequencies at loci responding to the very strong selection imposed by insect management practices. We conclude that this technology could be useful for identifying strong selection for resistance alleles across plant and insect species in agricultural ecosystems, providing an opportunity for detection and mitigation of widespread phenotypic resistance to management practices.

### Data Availability

Scripts and configuration files used for genome assembly can be found at: https://github.com/mcadamme/Hv_Genome_Assembly_Draft1

Scripts used for population genomic analysis can be found at: https://github.com/mcadamme/FieldHv_Pop_Genomics

Raw sequence ddRAD-seq data have been deposited in the following Dryad digital data repository: doi:10.5061/dryad.4k40j

Our *H. virescens* draft 1 assembly (accession NWSH00000000) and associated read sequences have been deposited in the NCBI database under BioProject number PRJNA379496.

## Acknowledgements

Thanks to Dr. J. Schaff and Dr. D. Baltzegar of the North Carolina State University Genomic Sciences Lab for their insightful suggestions on ways to improve our methods. We also thank Dr. Keith Hopper for early discussions of our genome assembly. Gabrielle Beaudry, Emma Thompson, Kelsey Mckinney, Wilfred Wong, Sharyar Samir, and Xuechun Wang isolated gDNA from the moths used in this project. Nico Olegario assisted with the L1029H genotyping. Mr. S. Micinski, Dr. J. Lopez, and Dr. J. Westbrook collected the moths used in this project. R. Waples wrote one of the custom scripts used in our data pipeline. This project was supported by the Biotechnology Risk Assessment Program competitive grant numbers 2012-33522-19793 and 2016-33522-25640 from the USDA - National Institute of Food and Agriculture.

