## Supplementary Materials for "Contemporary evolution of a Lepidopteran species, *Heliothis virescens*, in response to modern agricultural practices" (from another version of this preprint: version 3, 2017-07-30)

**Supporting Information**

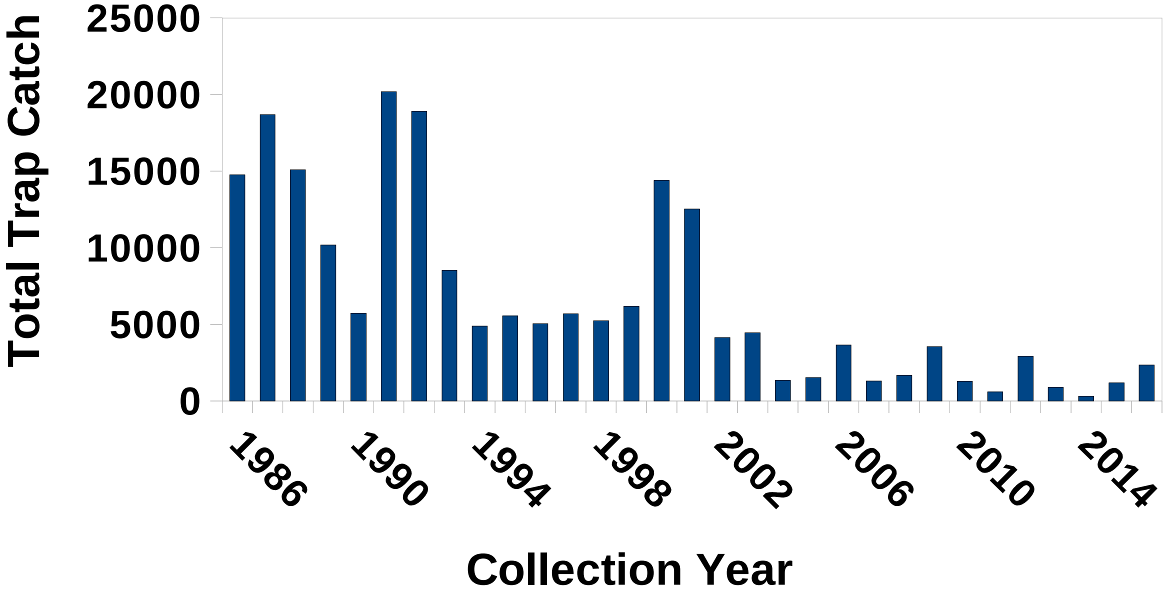


**Supplementary Figure 1 –** Total annual pheromone-baited trap catch of tobacco budworm from Bossier Parrish, LA, in the years 1986 through 2016.

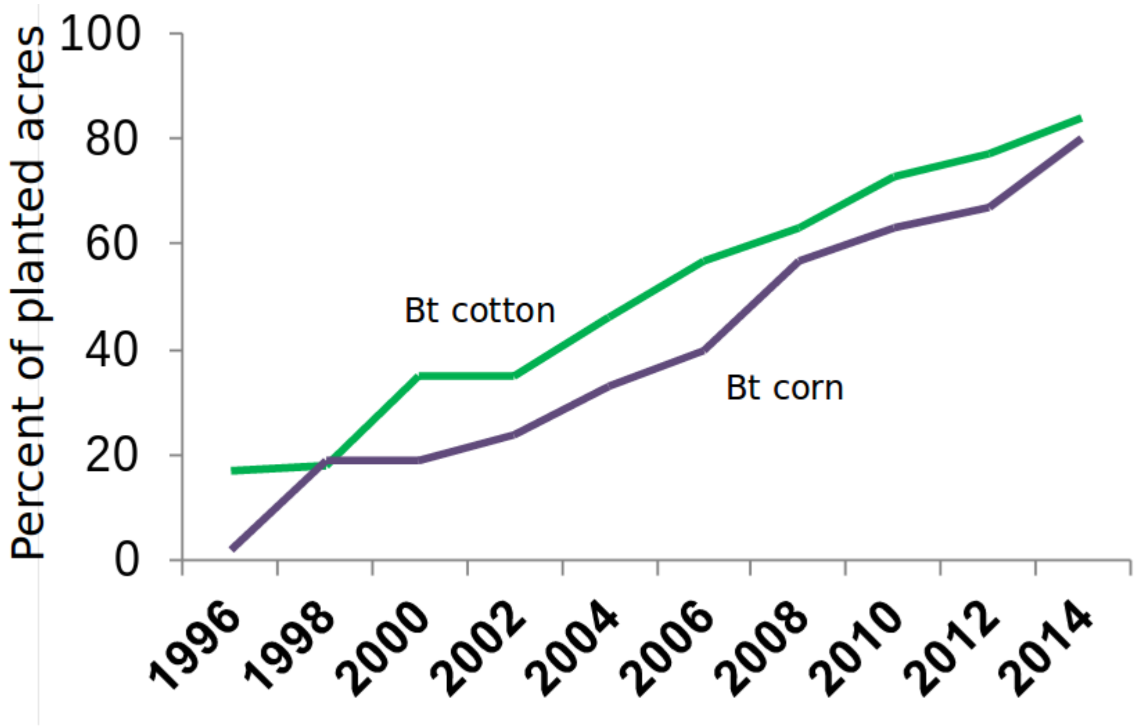


**Supplementary Figure 2 –** Adoption of transgenic cotton (green) and corn (blue) that expresses Bt toxins following their commercialization in the US in 1996. Transgenic crop adoption is expressed as a percentage of the overall planted acreage of each crop. Adapted from James 2015, USDA-ERS data.

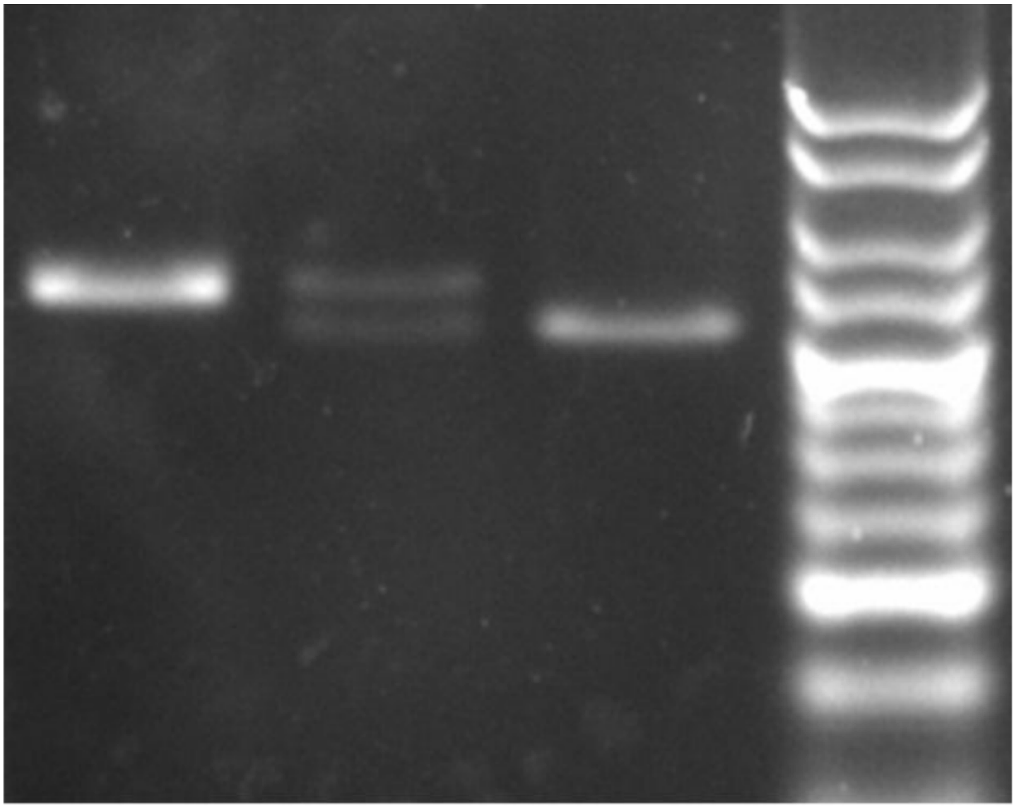

**Supplementary Figure 3 -** Three *H. virescens* individuals from Louisiana in 2012 were genotyped using Sanger sequencing. Amplicons from these same samples were digested with NlaIII and electrophoresed on a 3.5% agarose gel alongside Hyperladder V. From left to right, the genotypes as viewed on the agarose gel confirmed the Sanger genotypes, TT, AT, and AA.

```
Hv_11322_hap1     1  AATTCAATTAATTAAAAT----AATGTTGTTATGACATATGATAAGACTCAAACAGGTTT
Hv_11322_hap2     1  AATTCAATTAATTAAAAT----AATGTTGTTATGACATATGATAAGACTCAAACAGGTTT
Hv_11322_hap3     1  AATTCAATTAATTAAAATAATTAATGTTGTTATGACACATGATAAAACTCAAACAGGTTT
Hv_11322_hap4     1  AATTCAATTAATTAAAAT----AATGTTGTTATGACATATGATAAAACTCAAACAGGTTT
Hv_11322_hap5     1  AATTCAATTAATTAAAAT----AATGTTGTTATGACATATGATAAAACTCAAACAGGTTT

Hv_11322_hap1    57  GAAACGTGATTAAGAAAATGTTGTGTTCGATTTC--TTGTCAAATGCATGAATAAAAAAA
Hv_11322_hap2    57  GAAACGTGATTAAGAAAATGTTGTGTTCGATTTC--TTGTCAAATGCATGAATAAAAAAA
Hv_11322_hap3    61  GAAACGTGATTAAGAATATGTTGTGTTCGATTTT--TTGTCAAATGCATGAATA----AA
Hv_11322_hap4    57  GAAACGTGATTAAGAAAATGTTGTGTAAGATTTATTTTCTCAAACGCTTGAC-A----AA
Hv_11322_hap5    57  GAAACGTGATTAAGAGAATGTTGTGTCCGATTTTTTTCTCAAACGCATGAC-A----AA

Hv_11322_hap1   115  AACCTGTTTAACTTTTTCTAATAACACTT-ATAATATTT-TTAAACGA-----AAGTTTT
Hv_11322_hap2   115  AACCTGTTTAACTTTTTCTAATAACACTT-ATAATATTT-TTAAACGA-----AAGTTTT
Hv_11322_hap3   115  AACCTGTTTAGCTTTTTCTAATAACACTT-ATAATATTT-TTAAACGAAATAGAAGTTTT
Hv_11322_hap4   112  CATCTATTTAACTTCTTCTAATAACACTTAATAATATTTTTTAAACGAAATAGAAGTTTT
Hv_11322_hap5   112  CAT---CTTAACTTCTTCTAATAACACTT-ATAATATTTTTTAATCGAGAGAGAAGTTTT

Hv_11322_hap1   168  TGTAGCAGGAATAT-----ATGTACCTTAATTTTCCATTTGGAAAATAGGCAATCTTTAT
Hv_11322_hap2   168  TGTAGCAGGAATAT-----ATGTACCTTAATTTTCCATTTGGAAAATAGGCAATCTTTAT
Hv_11322_hap3   173  TGTAGCAGGAATAT-----ATGTACCTTAATTTTCCATTTGGAAAATAGGCAAACTTTAT
Hv_11322_hap4   172  TGTAGCAGAAATATATGCTATGTACCTTAATTTTCCATTTTGAAAATAGGCAAACTTT--
Hv_11322_hap5   168  TGTAGCAGAAATATATGCTATGTACCTTTATTTTCCATTTTGAAAATAGGCAAACTTT--

Hv_11322_hap1   223  TTATTAACTTTTTATTAGTACAGGCAGTCTGAAGCCGCGGCCCACAAATACGGGTCTCGT
Hv_11322_hap2   223  TTATTAACTTTTTATTAATACAGGCAGTCTGAAGCCGCGGCCCACAAATACGGGTCTCGT
Hv_11322_hap3   228  TTATTAACTTTTTATTAGTGCAGGCAGTCTGAAGTCGCGGCCCACAAATACGGGTCTCGT
Hv_11322_hap4   230  --ATTAACTTTTGATTAATACAGGCAGTCTGAAGTCGCGGCCCACAAATACGGGTCTCGT
Hv_11322_hap5   226  --ATTAACTTTTTATTAATACAGGCAGTCTGAAGTCGCGGCCCACAAATACGGGTCTCGT

Hv_11322_hap1   283  TCACGCACAATCAGGTCAAGGCATTTGTTTCACAGTCAAGTATCGCTGCAGCTTACTTTA
Hv_11322_hap2   283  TCACGCACAATCAGGTCAAGGCATTTGTTTCACAGTCAAGTATCGCTGCAGCTTGCTTTA
Hv_11322_hap3   288  TCACGCACAATCAGGTCAAGGCATTTGTTTCACAGTCAAGTATCGCTGCAGCTTGCTTTA
Hv_11322_hap4   288  TCACGCACAATCAGGTCAAGGCATTTGTTTCACAGTCAAGTATCGCTGCAGCTTACTTTA
Hv_11322_hap5   284  TCACGCACAATCAGGTCAAGGCATTTGTTTCACAGTCAAGTATCGCTGCAGCTTACTTTA

Hv_11322_hap1   343  ATACCGTT
Hv_11322_hap2   343  ATACAGTT
Hv_11322_hap3   348  AT-----A
Hv_11322_hap4   348  AT-----A
Hv_11322_hap5   344  ATACCG-T
```

**Supplementary Figure 4** - Multiple sequence alignment for the top 5 ddRAD-seq haplotypes found upstream from the *H. virescens* pyrethroid resistance locus. Hv_11322_hap1 appeared to be in strong linkage disequilibrium with voltage-gated sodium channel mutation L1029H.

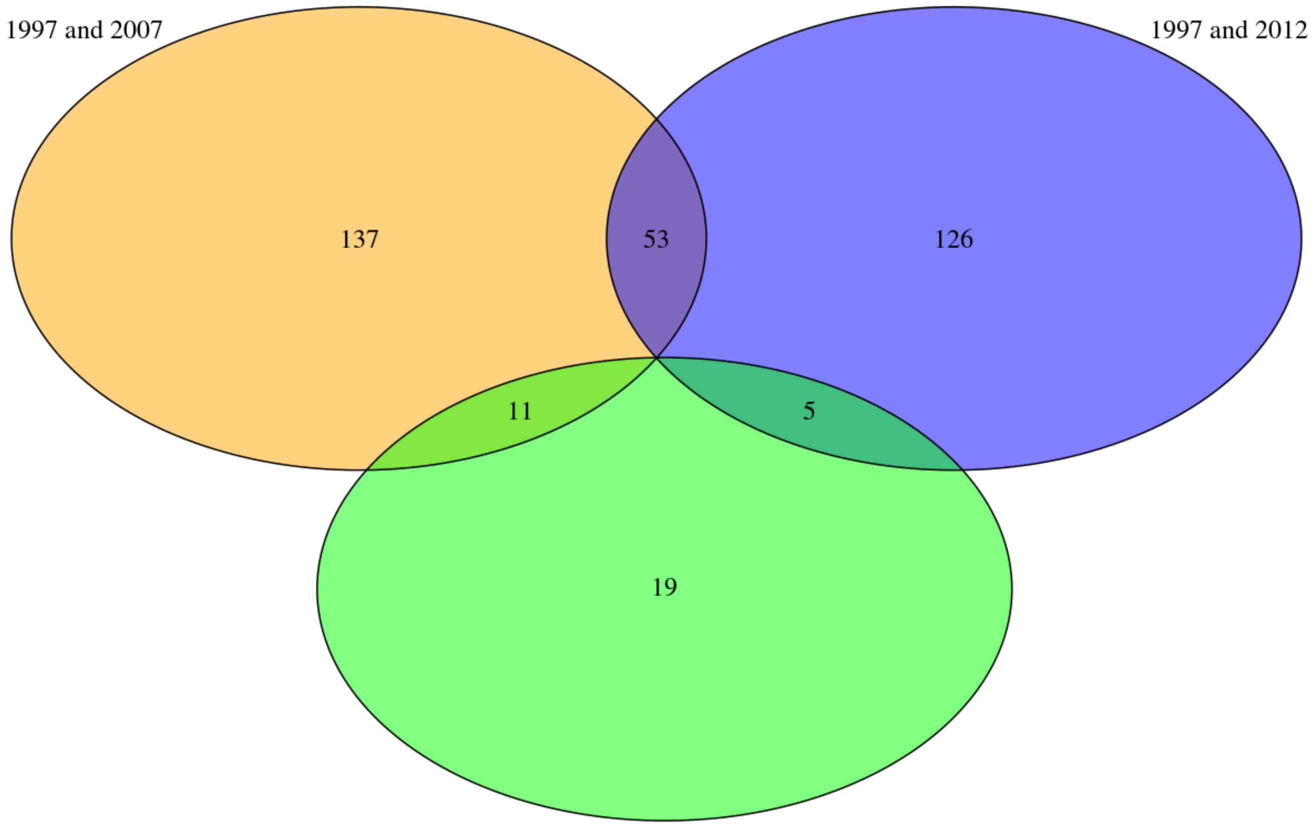


**Supplementary Figure 5 –** Venn diagram of the number of SNP outliers (of 8,963 SNPs) discovered through population genomic comparison between pairs of years for collections made in 1997, 2007 and 2012. Each by-year comparison is represented by a single circle, where the number of outliers is represented by the sum of the numbers within a single circle. Numbers found in the overlap between circles represent the number of SNPs that are shared outliers in more than one by-year comparison.

**Supplementary Table 1 –** GPS coordinates for pheromone-baited trap locations in Bossier Parrish, LA and Burleson County, TX.  All males collected from pheromone-baited traps within each state were held together in bottles by collection date.

| **State** | **Trap No.** | **Latitude (deg N)** | **Longitude (deg W)** |
|---|---|---|---|
| TX | Hv_101 | 30° 33' 22.572" | 96° 27' 57.096" |
| TX | Hv_103 | 30° 34' 40.728" | 96° 29' 27.06" |
| TX | Hv_303 | 30° 30' 45.36" | 96° 24' 20.592" |
| LA | Hv_20 | 32° 25' 7.66" | 93° 38' 5.03" |
| LA | Hv_21 | 32° 24' 59.55" | 93° 38' 18.62" |
| LA | Hv_24 | 32° 24' 51.28" | 93° 38' 7.05" |
| LA | Hv_26 | 32° 24' 50.77" | 93° 38 '30.84" |
| LA | Hv_28 | 32° 25' 19.03" | 93° 38 '30.85" |

**Supplementary Table 2 –** Genomes and partial sequences that were used to check for potential microbial/viral contamination, as well as filter out *H. virescens* mitochondrial reads.

| Species | Host | GenBank Accession |
|---|---|---|
| Phi-X | - | NC_001422.1 |
| *Wolbachia pipientis* | *D. melanogaster* | AE017196.1 |
| *Wolbachia pipientis* | *D. simulans* | CP003884.1 |
| *Wolbachia pipientis* | *C. lectularis* | AP013028.1 |
| Nuclear Polyhedrosis Virus | *H. zea* | NC_003349.1 |
| Ascovirus | *H. virescens* | JX491653.1 |
| *H. virescens* partial mtDNA | - | Sequence derived from a preliminary *H. virescens* genome assembly, followed by alignment to the *Bombyx mori* complete mtDNA genome. See Supplementary Data File 1. |

**Supplementary Table 3 –** *H. virescens* linkage groups for which mapped markers could be aligned to genome scaffolds. Scaffolds underlined and in bold aligned to multiple markers, where each belonged to different linkage groups. *B. mori* chromosome numbers that correspond to *H. virescens* LG were identified by Blastn alignment of mapped markers to the *B. mori* genome (Fritz *et al.* 2016).

| LG | *B. mori* Chr | Number of Scaffolds Aligned per LG | Scaffold Numbers |
|---|---|---|---|
| 1 | 25 | 21 | 3526, 432, 928, 169, 2746, 484, 668, 978, 5188, 635, 382, 7799, 799, 4346, 3173, 4198, 208, 90, 491, 438, 2041 |
| 2 | 8 | 16 | 3847, 73, 5371, 321, 6718, 7267, 1262, 2932, 591, 1247, 7011, 8227, 2333, 259, 3186, 8724 |
| 3 | 17 | 20 | 1440, 747, 702, 5972, 5688, 74, 32, 920, 428, 568, 4501, 2272, 3935, 226, 2376, 4667, 1912, 1259, 1428, 3835 |
| 4 | - | 7 | 45, 758, 2941, 65, 349, 708, 2481 |
| 5 | 4 | 26 | 12, 49, 589, 4491, 5502, 372, 80, 8105, 2928, 3200, 323, 818, 1209, 171, 1082, 10, 25, 1583, 1722, 330, 2986, 224, 192, 2132, 5931, 1957 |
| 6 | 5 | 32 | 131, 356, 284, 1804, 4231, 2886, 3791, 436, 2408, 629, 355, 1094, 616, 3360, 2529, 7892, 437, 1155, 4518, 299, 1052, 105, 1392, 469, 1823, 1115, 2453, 1104, 1446, 207, 3246, 1252 |
| 7 | 22 | 27 | 6059, 471, 5980, 8481, 921, 1463, 3345, 1153, 6174, 6209, 8502, 8293, 2822, 3694, 572, 2731, 2919, 4914, 2984, 3878, 8749, 2867, 3429, 271, 2123 |
| 8 | 11 | 20 | 1575, 3251, 1399, 4540, 294, 4139, 1019, 1944, 4110, 113, 600, 214, 1176, 2654, 617, 91, 796, 460, 619, 2636 |
| 9 | 23 | 6 | 180, 383, 5460, 3106, 415, 1873 |
| 10 | 3 | 15 | 2173, 1218, 793, 488, 1, 1347, 4374, 1835, 60, 2349, 3231, 122, 503, 2114, 3517 |

**Supplementary Table 3 continued.**

| LG | *B. mori* Chr | Number of Scaffolds Aligned per LG | Scaffold Numbers |
|---|---|---|---|
| 11 | 9 | 22 | 1926, 5077, 202, 2724, 7924, 5135, 57, 1768, 483, 2105, 8497, 1853, 2363, 1690, 2258, 36, 7674, 8541, 1365, 4097, 7816, 3696 |
| 12 | - | 15 | 1740, 2286, 2269, 243, 3697, 1368, 752, 789, 3524, 2436, 3161, 43, 3333, 4344, 7880 |
| 13 | 13 | 23 | 6089, 2426, 3463, 834, 1653, **209**, 6485, 2410, 1732, 370, 1480, 3879, 546, 188, 378, 723, 1540, 2281, 2661, 1301, 374, 2467, 6005 |
| 14 | 19 | 16 | 15, 622, 1015, 7200, 1895, 7088, 822, 7284, 410, 1126, 1144, 1932, 3618, 84, 2577, 2205 |
| 15 | 6 | 21 | 3522, 430, 223, 531, 2977, 408, 1855, 6810, 283, 149, 849, 7955, 2789, 2605, 1062, 1881, 1372, 1045, 684, 151, 3140 |
| 16 | 23 | 17 | 1090, 4593, 982, 19, 933, 1220, 2328, 6931, 422, 1374, 1393, 56, 3262, 1861, 3868, 1669, 8809 |
| 17 | 7 | 11 | 1191, 784, 304, 1755, 2127, 197, 1138, 3092, 2608, 2495, 2416 |
| 18 | 26 | 10 | 1645, 2624, 8, 6278, 2081, 7301, 2739, 6494, 1160, 4298 |
| 19 | 11, *22* | 2 | 398, 872 |
| 20 | 10 | 18 | 314, 729, 0, 4397, 4138, 3610, 750, 3452, 2032, 286, 2186, 129, 7383, 13, 886, 1012, 942, 5764 |
| 21 | 21 | 20 | 1064, 1050, 1693, 1363, 3343, 1571, 3448, 884, 1921, 1040, 860, 2040, 586, 5202, 716, 92, 210, 579, 877, 2052 |
| 22 | 15 | 24 | 637, 549, 291, 4766, 2070, 2159, 1760, 836, 599, 301, 176, 516, 5965, 739, 1282, 2149, 247, 352, 114, 4558, **121**, 86, 11, 95 |
| 23 | 14 | 25 | 463, 4166, 1982, 380, 218, 2737, 508, 1782, 4183, 870, 4475, 903, 5148, 709, 749, 5350, 3714 |
| 24 | - | 5 | 1735, 1817, 3950, **1860**, 3199 |
| 25 | 12, *1* | 36 | 8241, 3469, 1651, 847, 29, 625, 2134, 297, 1194, 2639, 512, 1486, **121**, 735, 119, 8357, 780, 4596, 1748, 2199, 1518, 303, 2047, 1506, 1198, 1195, 3436, 240, 4440, 1629, 509, 8774, 309, 1344, 3016, 452 |
| 26 | - | 8 | 511, 772, 100, 4077, 3129, 101, 4, 1515 |
| 27 | 2 | 3 | 5329, 6212, 3949 |
| 28 | - | 4 | 1263, 83, 75, 580 |
| 29 | 16 | 6 | 155, 885, 3424, 3664, 1303, 63 |
| 30 | - | 6 | 363, 1178, 158, **209**, 477, 290 |
| 31 | - | 6 | 1799, 67 4649, 730, 792, 3045 |
| 32 | - | 2 | 1588, 4144 |
| 33 | 14, *28* | 6 | 394, 1877, 1710, **1860**, 419, 449 |

**Supplementary Table 4** – *H. virescens* genome assembly specifications.

| **Analysis** | **Assembly Results** |
|---|---|
| Total Genome Length | 403,154,421 bp |
| Number of Scaffolds | 8,826 |
| Number of Ns/Gaps | |
| Total | 38,034,845 bp |
| Number of Occurrences | 47,236 |
| Mean Size | 805 bp |
| Mean Scaffold Size | 45,678 bp |
| Scaffold Size Range | 659 – 628,964 bp |
| Scaffold N50 Size | 102,124 bp |
| Scaffold N90 Size | 21,868 bp |
| BUSCO (v. 2.0.1) | |
| Complete Core Genes | 865 (of 978) |
| Single Copy Complete | 812 |
| Duplicated Copy Complete | 53 |
| Fragmented | 39 |
| Missing | 74 |
| Number of Genes | 15,092 |
| Number of mRNAs | 18,739 |
| Number of Exons | 140,252 |
| Number of Introns | 121,513 |
| Number of CDS | 18,739 |
| Gene Length | |
| Shortest | 151 bp |
| Longest | 187,213 bp |
| Mean | 9,105 bp |
| CDS Length | |
| Shortest | 129 bp |
| Longest | 34,770 bp |
| Mean | 1615 bp |
| % Genome Covered by Genes | 34.1 |
| % Genome Covered by CDS | 7.5 |
| Mean mRNAs per gene | 1 |
| Mean exons per mRNA | 7 |
| Mean introns per mRNA | 6 |
